# Ancestral lineage of SARS-CoV-2 is more stable in human biological fluids than Alpha, Beta and Omicron variants of concern

**DOI:** 10.1101/2022.08.17.504362

**Authors:** Taeyong Kwon, Natasha N. Gaudreault, David A. Meekins, Chester D. McDowell, Konner Cool, Juergen A. Richt

## Abstract

SARS-CoV-2 is a zoonotic virus which was first identified in 2019, and has quickly spread worldwide. The virus is primarily transmitted through respiratory droplets from infected persons; however, the virus-laden excretions can contaminate surfaces which can serve as a potential source of infection. Since the beginning of the pandemic, SARS-CoV-2 has continued to evolve and accumulate mutations throughout its genome leading to the emergence of variants of concern (VOCs) which exhibit increased fitness, transmissibility, and/or virulence. However, the stability of SARS-CoV-2 VOCs in biological fluids has not been thoroughly investigated so far. The aim of this study was to determine and compare the stability of different SARS-CoV-2 strains in human biological fluids. Here, we demonstrate that the ancestral strain of Wuhan-like lineage A was more stable than the Alpha VOC B.1.1.7, and the Beta VOC B.1.351 strains in human liquid nasal mucus and sputum. In contrast, there was no difference in stability among the three strains in dried biological fluids. Furthermore, we also show that the Omicron VOC B.1.1.529 strain was less stable than the ancestral Wuhan-like strain in liquid nasal mucus. These studies provide insight into the effect of the molecular evolution of SARS-CoV-2 on environmental virus stability, which is important information for the development of countermeasures against SARS-CoV-2.

**Importance:** Genetic evolution of SARS-CoV-2 leads to the continuous emergence of novel variants, posing a significant concern to global public health. Five of these variants have been classified so far into variants of concern (VOCs); Alpha, Beta, Gamma, Delta, and Omicron. Previous studies investigated the stability of SARS-CoV-2 under various conditions, but there is a gap of knowledge on the survival of SARS-CoV-2 VOCs in human biological fluids which are clinically relevant. Here, we present evidence that Alpha, Beta, and Omicron VOCs were less stable than the ancestral Wuhan-like strain in human biological fluids. Our findings highlight the potential risk of contaminated human biological fluids in SARS-CoV-2 transmission and contribute to the development of countermeasures against SARS-CoV-2.

## Introduction

Severe acute respiratory syndrome coronavirus 2 (SARS-CoV-2) is a recently emerging respiratory virus and the causative agent for the current pandemic. SARS-CoV-2 belongs to the genus *Betacoronavirus*, family *Coronaviridae*, order *Nidovirales*; it is an enveloped virus containing a positive-sense, single-stranded RNA genome of approximately 30 kb in length. The first two thirds of the genome (5’ to 3’) are comprised of two overlapping open reading frames (ORF), ORF1a and ORF1b, which encode two large polyproteins through a programmed −1 ribosomal frameshifting (1). These polyproteins are proteolytically cleaved into a total of 16 active nonstructural proteins, which are involved in virus replication and transcription as well as innate immune evasion by suppressing host factors in various signaling pathways (2). The last third of the genome encodes four structural proteins: spike (S), envelope (E), membrane (M), and nucleocapsid (N), and eight accessory proteins: ORF3a, ORF3b, ORF6, ORF7a, ORF7b, ORF8, ORF9a and ORF9b. The trimeric S protein of SARS-CoV-2 mediates viral attachment to the host cell receptor, human angiotensin-converting enzyme 2 (ACE2), and subsequent virus-cell fusion and entry into target cells (3). The E and M proteins are components of the viral envelope and play a role in viral assembly and budding of SARS-CoV-2 (4-6). The N is an RNA-binding protein responsible for viral genome packaging (7).

SARS-CoV-2 has continued to evolve and has accumulated mutations throughout its genome since its emergence in late 2019. Due to the error-prone nature of RNA-dependent RNA polymerase, random mutations are introduced into the genome of SARS-CoV-2 during viral replication such that SARS-CoV-2 populations exist as a quasispecies. Selection pressures drive the viral population to maintain beneficial mutations which could potentially increase viral fitness. This process of mutation and selection guides the evolution of SARS-CoV-2 and contributes to the emergence and spread of new variants that can pose an increased risk to global public health. Five variants of SARS-CoV-2 have so far been designated as Variants of Concern (VOCs): Alpha (B.1.1.7), Beta (B.1.351), Gamma (P.1), Delta (B.1.617.2), and Omicron (B.1.1.529) (see: https://www.who.int/en/activities/tracking-SARS-CoV-2-variants). The Alpha VOC was first identified in the United Kingdom in September 2020 and became the dominant strain circulating in many parts of the world until May 2021 (8). The Beta and Gamma VOCs were first identified in South Africa and Brazil, respectively; they have been responsible for small proportions of COVID-19 cases worldwide, but their emergence and continued spread have been highlighted due to their ability to evade pre-existing immunity and therapeutics (9, 10). After being first identified in October 2020 in India, the Delta VOC has spread to many other countries and replaced the previously prevalent Alpha VOC to become the dominant strain of SARS-CoV-2 worldwide (11). On 26 November 2021, the variant B.1.1.529 was designated as the Omicron VOC, which is a highly divergent variant harboring a high number of mutations, especially in the S protein. In addition, many other variants with specific genetic makers which are predicted to affect virus characteristics have been classified as Variants of Interest (VOIs).

To date, antigenic and virological aspects of SARS-CoV-2 VOCs, such as transmissibility, disease severity, vaccine and therapeutic efficacy and immune evasion, have been widely investigated. Most studies have explored the stability of several strains of SARS-CoV-2 on the surfaces (12-14) and a few have investigated the stability of SARS-CoV-2 in human biological fluids (15-17). These studies enable us to determine the potential of SARS-Cov-2 transmission by fomites. However, the stability of different VOCs in human biological fluids has not been compared side-by-side. Studies evaluating the stability of new variant strains of SARS-CoV-2 in biological fluids is important for assessing the risk of potential fomite transmissions. Therefore, we first evaluated the stability of three different SARS-CoV-2 strains, (1) an ancestral lineage A strain, (2) an Alpha VOC and (3) a Beta VOC, in human biological fluids under different environmental conditions. In addition, the stability of the Omicron VOC BA.1 was assessed in biological fluids.

## Results

To characterize the detailed genetics of the virus stocks used in this study, we performed next generation sequencing (NGS) using an Illumina Nextseq platform. The results showed that the consensus sequence of the WA-1 strain was 100% identical with the reference sequence available in GISAID (accession ID: EPI_ISL_404895) except for a synonymous mutation from C to T at position 1912. The virus stock of the Alpha VOC was 100% homologous with the GISAID reference sequence (accession ID: EPI_ISL_683466), and had several amino acid substitutions in the S protein when compared to the WA-1 strain: H69del, V70del, Y145del, N501Y, A570D, D614G, P681H, T716I, S982A, and D1118H (Supplementary table 1). We also found an amino acid substitution from aspartic acid to leucine at position 3 of the nucleocapsid protein (D3L).. The virus stock of the Beta VOC was homolgous with the GISAID reference sequence (accession ID: EPI_ISL_678615) and contained several amino acid substitutions in the S protein when compared to the WA-1 strain: L18F, D80A, D215G, L242del, A243del, L242del, K417N, E484K, N501Y, D614G, and A701V. Two additional substitutions, Q677H and R682W, were found in the Beta VOC virus stock when compared to the reference sequence. Also, proline at position 71 of the E protein was replaced with leucine in the Beta VOC (P71L) and a P252L substitution was found in NSP5 and the substitution R115L in ORF8. The consensus nucleotide sequence of the Omicron VOC stock had 100% identity to the reference sequence deposited in GISAID (accession ID: EPI_ISL_7908052). In the Omicron VOC, a total 31 amino acid substitutions, six deletions, and three insertions were found in the S protein as well as several substitutions and deletions in other three structural proteins, when compared to the WA-1 strain.

In liquid nasal mucus (Fig. 1), the WA-1 strain was more stable than the Alpha VOC under indoor, summer, and spring/fall conditions (*p* < 0.0001). Similarly, the Beta VOC survived longer than the Alpha VOC under indoor (*p* < 0.001), summer (*p* < 0.0001), and spring/fall (*p* < 0.0001) conditions. In addition, we found a significant difference in the viral decay rate between the WA-1 strain and the Beta VOC under indoor conditions (*p* < 0.01). In winter conditions, the WA-1 strain and the Alpha VOC survived significantly longer than the Beta VOC (*p* < 0.05 for WA-1 vs Beta VOC; and *p* < 0.01 for Alpha VOC vs Beta VOC).

**Fig. 1.**
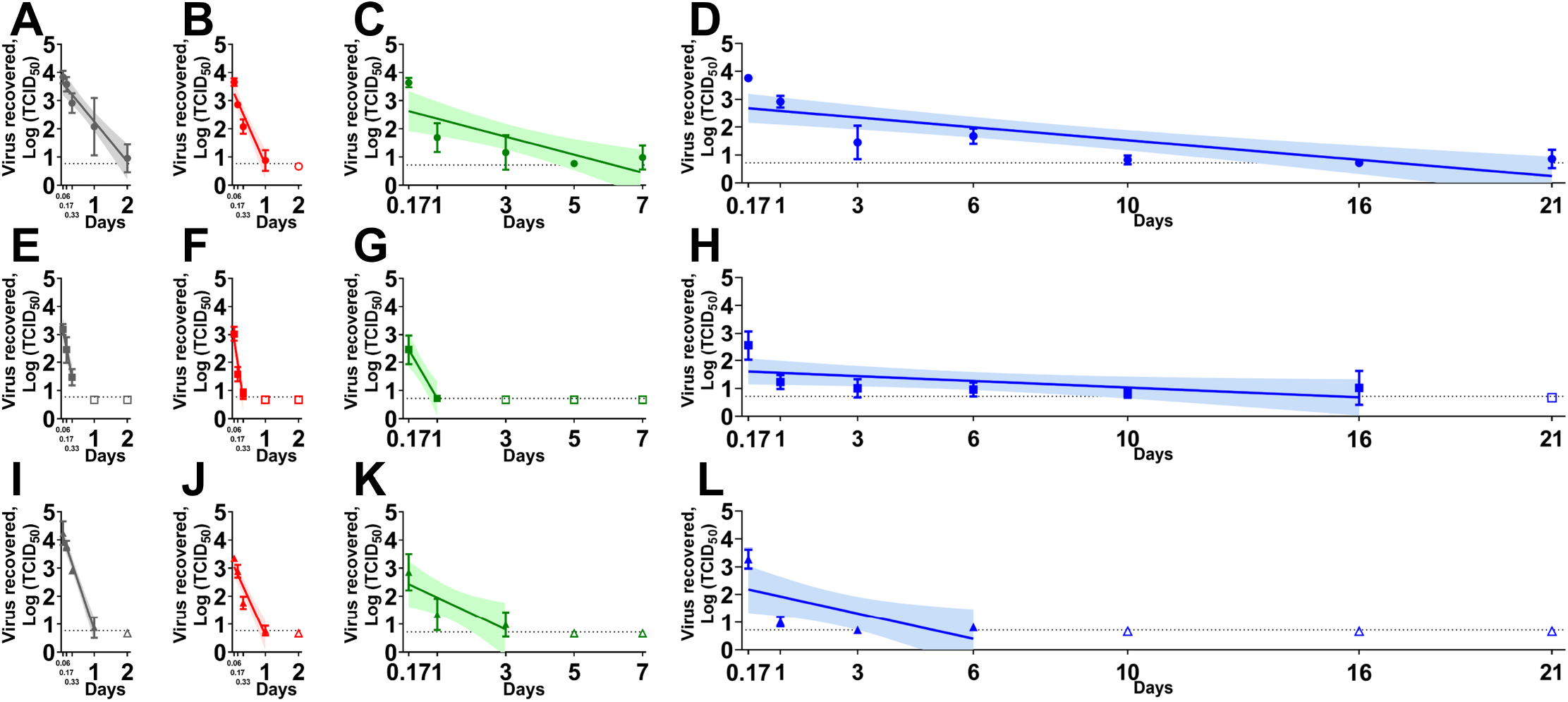
The stability of the severe acute respiratory syndrome coronavirus 2 (SARS-CoV-2) ancestral lineage A strain (A–D), the Alpha variants of concern (VOC) (E–H), and the Beta VOC (I–L) in human nasal mucus under indoor (A, E and I), summer (B, F and J), spring/fall (C, G and K), and winter (D, H and L) conditions. The cell culture derived virus (5 × 10^4^ TCID_50_) was mixed with nasal mucus in a 2 mL sealed tube and placed in a temperature and humidity-controlled chamber. After the incubation under each environmental condition, the sample was diluted in the 2 mL medium, filtered through 0.45 μm syringe filter, and titrated on Vero-TMPRSS2 cells. Virus titers were log-transformed to estimate a simple linear regression model. Virus titer at each time point was expressed as a geometric mean of three replicates and the standard deviation. A best-fit line and its 95% confidence interval of each regression model are represented by a solid line and its shade area. The dashed line indicates the limit of detection where at least one sample out of the three replicates was positive by virus isolation, and the empty symbols represent negative samples in all three replicates. On the *x*-axis, 0.06, 0.17 and 0.33 days are equal to 1.5, 4, and 8 hours, respectively.

In liquid sputum (Fig. 2), the WA-1 strain survived significantly longer than the Alpha VOC under indoor (*p* < 0.05), summer (*p* < 0.0001), spring/fall (*p* < 0.0001), and winter (*p* < 0.01) conditions. In addition, the WA-1 strain was also more stable than the Beta VOC in liquid sputum under indoor (*p* < 0.01), summer (*p* < 0.05), spring/fall (*p* < 0.0001), and winter (*p* < 0.01) conditions. Overall, the Alpha and Beta VOCs had similar stability in sputum under all conditions tested.

**Fig. 2.**
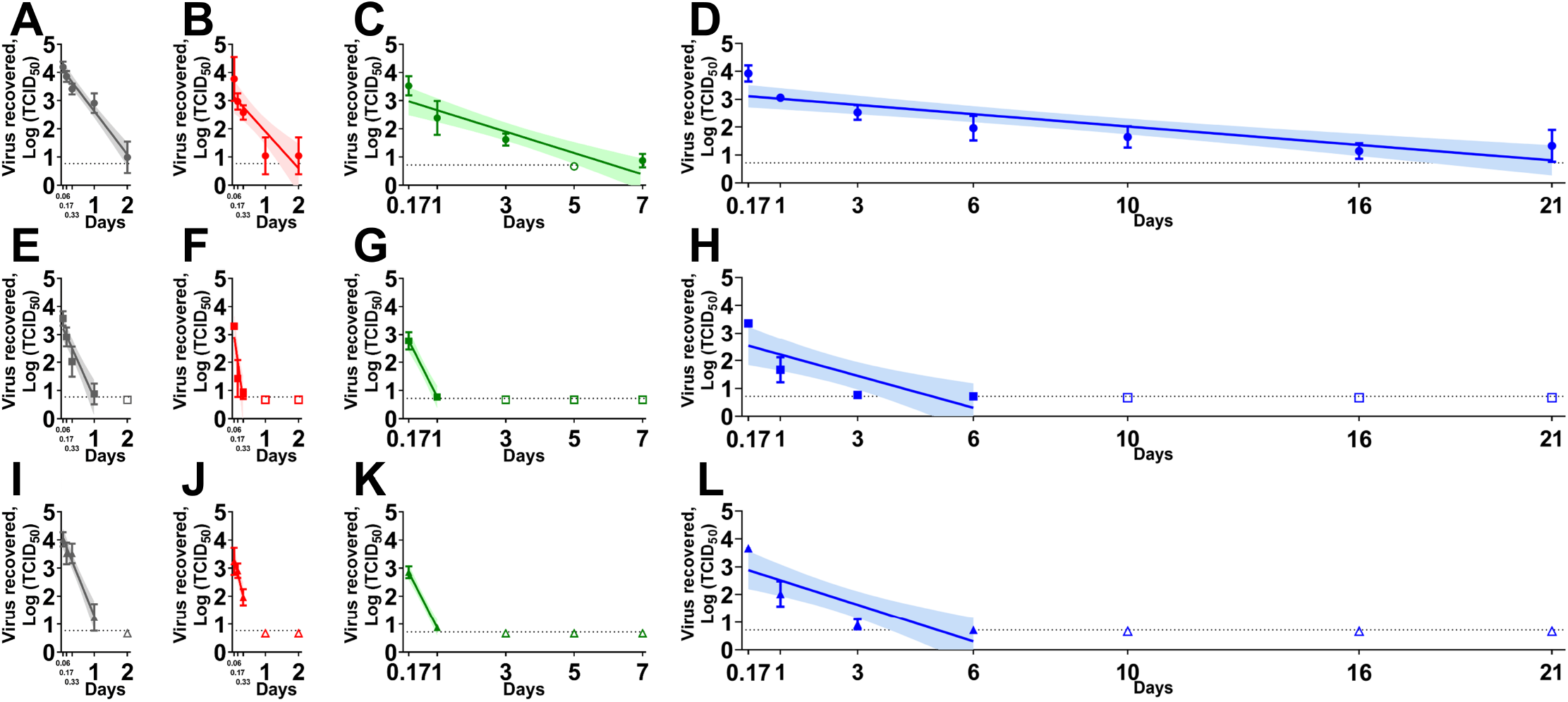
The stability of the severe acute respiratory syndrome coronavirus 2 (SARS-CoV-2) ancestral lineage A strain (A–D), the Alpha variants of concern (VOC) (E–H), and the Beta VOC (I–L) in human sputum under indoor (A, E and I), summer (B, F and J), spring/fall (C, G and K), and winter (D, H and L) conditions. The cell culture derived virus (5 × 10^4^ TCID_50_) was mixed with sputum in a 2 mL sealed tube and placed in a temperature and humidity-controlled chamber. After the incubation under each environmental condition, the sample was diluted in the 2 mL medium, filtered through 0.45 μm syringe filter, and titrated on Vero-TMPRSS2 cells. Virus titers were log-transformed to estimate a simple linear regression model. Virus titer at each time point was expressed as a geometric mean of three replicates and the standard deviation. A best-fit line and its 95% confidence interval of each regression model are represented by a solid line and its shade area. The dashed line indicates the limit of detection where at least one sample out of the three replicates was positive by virus isolation and the empty symbols represent negative samples in all three replicates. On the *x*-axis, 0.06, 0.17 and 0.33 days are equal to 1.5, 4, and 8 hours, respectively.

In liquid saliva (Table 1), the WA-1 strain and the Alpha VOC were more stable than the Beta VOC under winter conditions (*p* < 0.001 for WA-1 vs Beta VOC; and *p* < 0.01 for Alpha VOC vs Beta VOC). In liquid medium (Table 1), the half-life values of WA-1 were significantly higher than those of both, the Alpha and Beta VOCs under winter conditions (*p* < 0.05 for WA-1 vs Alpha VOC; and *p* < 0.001 for WA-1 vs Beta VOC). However, no difference in half-life values among the three strains was found in medium, nasal mucus, sputum, and saliva dried on the stainless steel surface (Table 1).

**Table 1.**
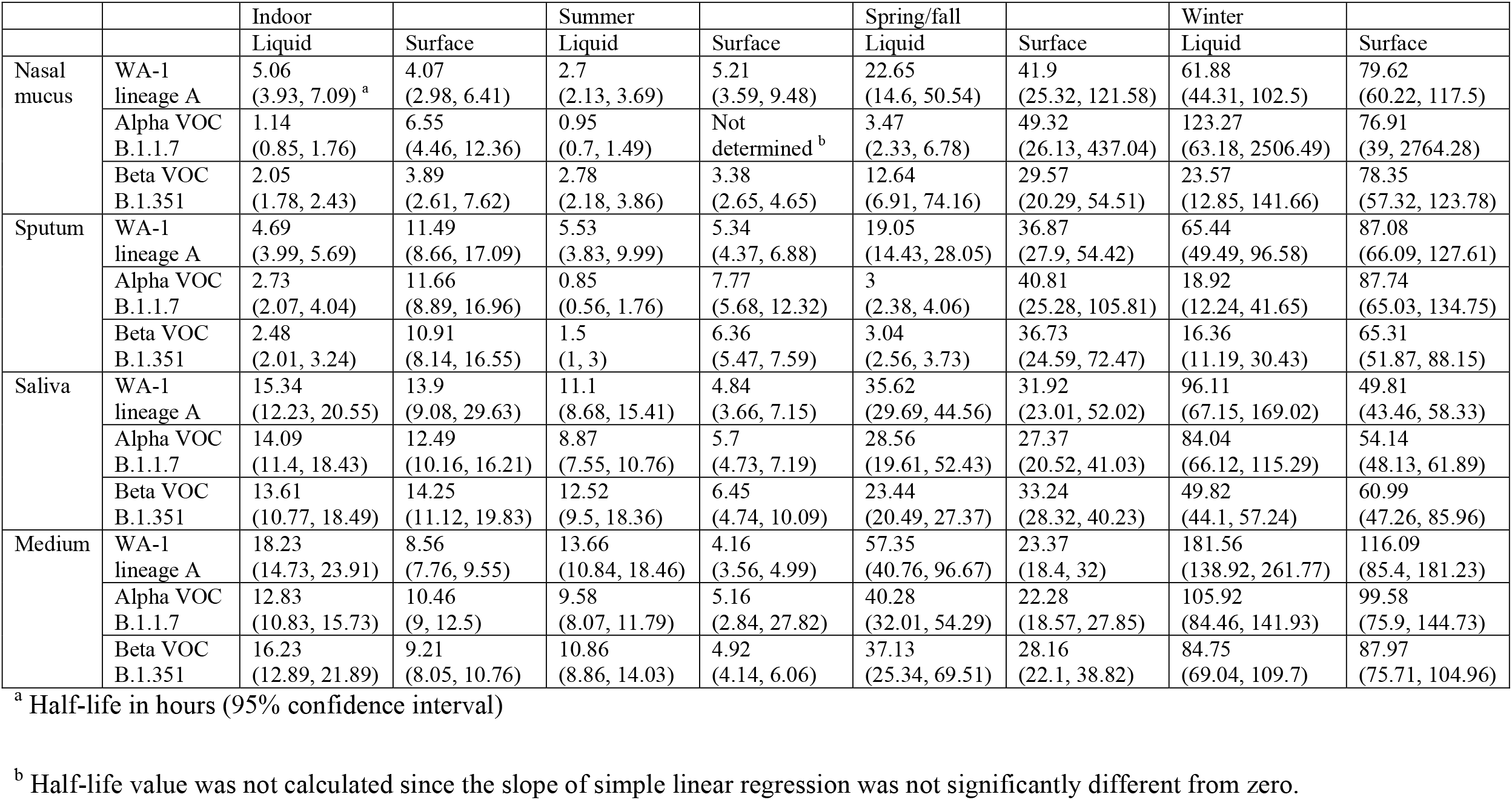
Half-life values of the SARS-CoV-2 ancestral lineage A strain and two variants of concern in liquid biological fluids or dried on surface under indoor, summer, spring/fall, and winter conditions.

In addition, the Omicron VOC was less stable than the WA-1 strain in liquid nasal mucus (*p* < 0.01) and medium (*p* < 0.01) under spring/fall conditions (Fig. 3). However, a significant difference was not observed in liquid sputum (Table 2).

**Fig. 3.**
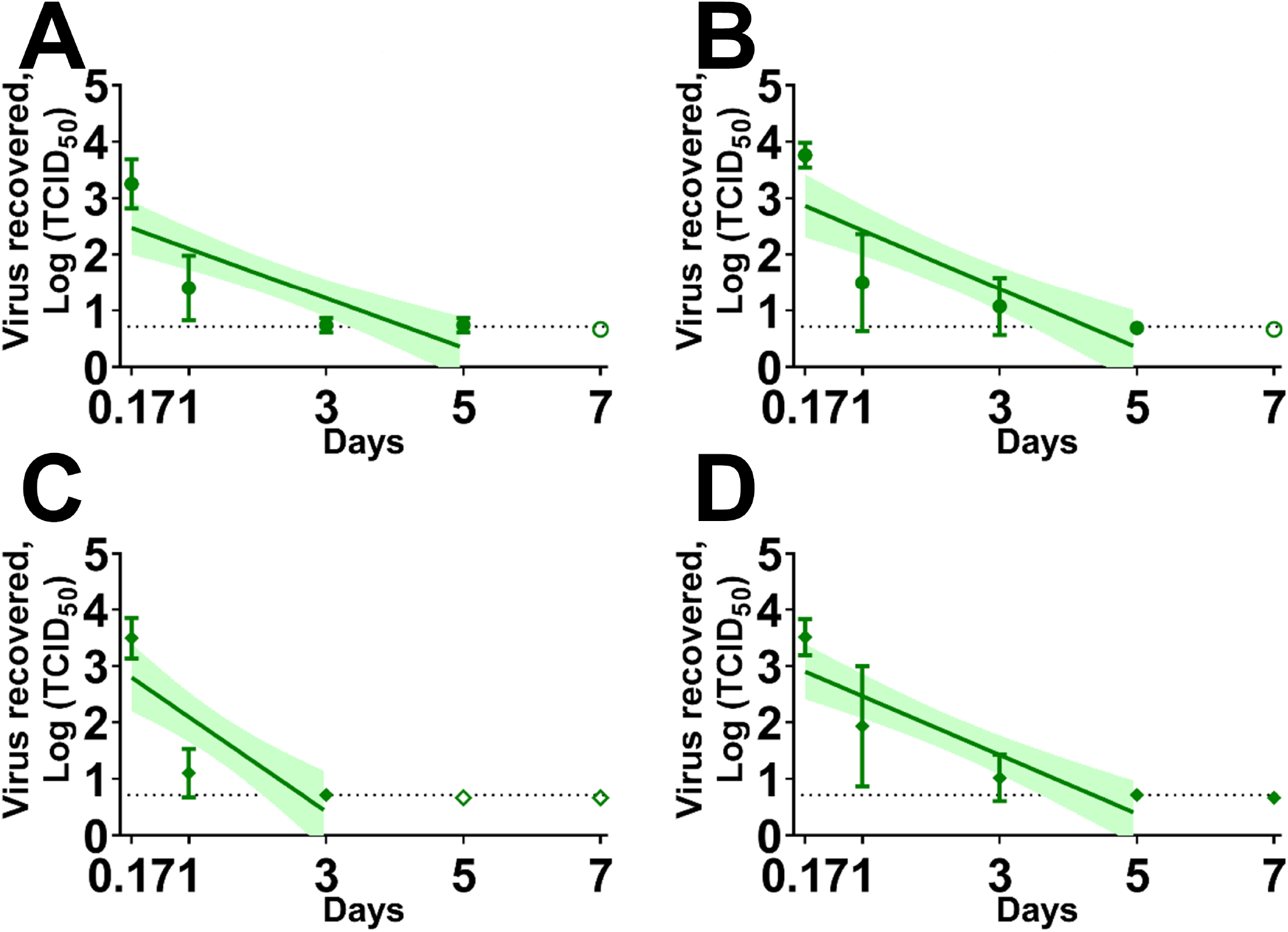
The stability of the severe acute respiratory syndrome coronavirus 2 (SARS-CoV-2) ancestral lineage A strain (A and B) and the Omicron variants of concern (VOC) (C and D) in human nasal mucus (A and C) and sputum (B and D) spring/fall conditions. The cell culture derived virus (3.1 × 10^4^ TCID_50_) was mixed with nasal mucus or sputum in a 2 mL sealed tube and placed in a temperature and humidity-controlled chamber. After the incubation, the sample was diluted in the 2 mL medium, filtered through 0.45 μm syringe filter, and titrated on Vero-TMPRSS2 cells. Virus titers were log-transformed to estimate a simple linear regression model. Virus titer at each time point was expressed as a geometric mean of three replicates and the standard deviation. A best-fit line and its 95% confidence interval of each regression model are represented by a solid line and its shade area. The dashed line indicates the limit of detection where at least one sample out of the three replicates was positive by virus isolation and the empty symbols represent negative samples in all three replicates. On the *x*-axis, 0.17 day are equal to 4 hours.

**Table 2.** Half-life values of the SARS-CoV-2 Omicron variants of concern in liquid nasal mucus and sputum under spring/fall conditions.

|  |  |  |
| --- | --- | --- |
|  |  | Spring/fall |
|  |  | Liquid |
| Nasal mucus | WA-1 lineage A | 16.4<br>(11.93, 26.2) <sup>a</sup> |
|  | Omicron VOC B.1.1.529 | 8.64<br>(6.1, 14.81) |
| Sputum | WA-1 lineage A | 13.89<br>(10.09, 22.3) |
|  | Omicron VOC B.1.1.529 | 13.89<br>(10.44, 20.75) |
| Medium | WA-1 lineage A | 71.38<br>(45.37, 167.24) |
|  | Omicron VOC B.1.1.529 | 30.32<br>(22.6, 46.04) |

<sup>a</sup> Half-life in hours (95% confidence interval)

## Discussion

SARS-CoV-2 can be excreted in many types of biological fluids from infected individuals, but nasal mucus, sputum, and saliva are the primary components in the generation of respiratory droplets that play a major role in SARS-CoV-2 transmission (18). The infectious droplets can be inhaled directly, which is a primary route of SARS-CoV-2 transmission. In addition, very fine droplets can evaporate quickly in the air to create infectious aerosols, which can remain infectious for hours (13). In contrast, the larger droplets drop down to the nearby area within a few minutes and contaminate surfaces with infectious virus (see: https://www.cdc.gov/coronavirus/2019-ncov/science/science-briefs/sars-cov-2-transmission.html). The virus that survives on surfaces can then be transferred by hand or other means to mucosal membranes in the oral or nasal cavity from contaminated surfaces (fomite transmission). Given this scenario, SARS-CoV-2 survival is of concern for its potential role in fomite transmission; this has led to the extensive investigations on the effect of surfaces (13, 19, 20), substrates (15-17, 21), and environmental factors (12, 22, 23) on SARS-CoV-2 stability.

It has been considered that virus stability outside the host is dependent on the intrinsic nature of the virus, type of surface, surrounding substrate, and environmental conditions. Previously, we reported extended SARS-CoV-2 survival under winter conditions compared to other seasonal climates on various surfaces and biological fluids, and differential virus decay rates depending on the types of surfaces and the type of human biological fluid (15, 20). In this study, to elucidate the effect of virus strain on virus stability, we prepared virus stocks of four SARS-CoV-2 strains using identical cell culture conditions to eliminate other variables besides the genetic of the virus. SARS-CoV-2 has an approximately 30 kb RNA genome encapsulated by the nucleocapsid protein, and the RNA-nucleocapsid complex is surrounded by an envelope made of a lipid bilayer from host cells. The spike, envelope, and membrane proteins are embedded in this outer lipid membrane and determine the distinct shape and structure of SARS-CoV-2. It is reasonable to suggest that these structural factors, such as structural proteins and lipid membrane, are primarily related to virus stability of individual SARS-CoV-2 strains. In this context, amino acid differences in the structural proteins may play a critical role in differential virus stability as described in this study.

A well-characterized SARS-CoV-2 mutation is the aspartic acid to glycine substitution at the position 614 of the spike protein (D614G). This substitution was found in early 2020 in Europe and quickly became dominant worldwide. Initial observations showed that patients infected with the D614G variant had higher viral loads with a significant effect on SARS-CoV-2 infectivity and transmissibility (24). Further studies demonstrated that SARS-CoV-2 harboring the single D614G mutation exhibited increased competitive fitness, and rapid transmission in primary human respiratory cells and animal models (25). Subsequent cryo-EM analysis indicated that the D614G substitution abolishes an inter-protomer salt bridge between D614 and K854 and/or an inter-protomer hydrogen bond between D614 and T859 which results in structural flexibility of the spike protein, allowing the S1 subunit of the protomer to be easily disassociated from the S2 subunit of the adjacent protomer (26-28). The resulting destabilization of the S1-S2 interface triggers a conformational change of the receptor binding domain (RBD) of the spike protein, toward the “up state”, leading to increased ACE2 binding and enhanced infectivity. All VOCs used in this study harbor this potentially destabilizing D614G substitution within the spike protein which could explain the reduced stability compared to the ancestral strain containing the D614. Another common mutation is N501Y in the spike protein, which is located within the receptor binding motif. This substitution influences the ability to bind ACE2 and evade antibody neutralization, however it does not result in notable structural changes (29-31). Furthermore, for the Alpha VOC, the A570D substitution in the S1 subdomain of the spike protein forms a new salt bridge with K854 in the RBD “up state” conformation or K964 in RBD “down state” conformation (30, 31), or it forms a new hydrogen bond with N856 (32). In contrast, the S982A substitution in the S2 subdomain of the spike protein causes the loss of a hydrogen bond with G545 (30) or T547 (32). In addition, the D1118H substitution in the S2 subdomain of the spike protein forms a symmetric histidine triad near the base of the spike, whereas the T716 substitution eliminates an intra-protomer hydrogen bond with Q1071 (32). For the Beta VOC, the E484K substitution in the S1 subdomain of the spike protein results in the elimination of a hydrogen bond with F490, and destabilizes the RBD structure with a high frequency of disordered RBDs (32). The overall architecture of the Omicron VOC spike is similar to those of other SARS-CoV-2 strains, but several of its substitutions introduce an inter-protomer new hydrogen bond or salt bridge between N856K and D568/T572 and a hydrogen bond between N764K and T315 (33, 34). In addition, other mutations in the nucleocapsid and envelope proteins of the VOCs might have effects on virus stability, although their roles in the structural stability of the virus particle remains unclear. It is likely that all amino acid substitutions which stabilize or destabilize the structure of the spike protein, as well as potentially other structural proteins, synchronously affect the stability of SARS-CoV-2.

We found significant differences among the stability of SARS-CoV-2 strains in various liquid biological fluids and medium, whereas no difference was observed in dried medium and biological fluids on a stainless steel surface. In particular, the WA-1 strain and the Beta VOC were more stable than the Alpha VOC under indoor, summer, spring/fall conditions in human nasal mucus, and the WA-1 strain was more stable than both VOCs under all four conditions in human sputum. Furthermore, the WA-1 strain was more stable than the Omicron VOC in nasal mucus. These results might imply that components such as certain enzymes present in the liquid biological fluids are responsible for different virus decay rates, since the drying process on surfaces causes the inactivation of these components due to lack of water. It is plausible that an active component in biological fluids exerts its antiviral activity more effectively against VOCs that harbor structurally less stable spike proteins in their envelopes. Previous studies have indicated that the single D614G substitution increased the stability of SARS-CoV-2. Planta *et al*. showed that the G614 virus was more stable than the D614 in Dulbecco’s PBS at 33°C, 37°C, and 42°C (35). Another study demonstrated that the D614 virus in DMEM lost a considerable degree of infectivity with a 3-log reduction between 14 and 30 days at 4°C, whereas only a 1-log reduction was found with the G614 virus (36). Moreover, two recent studies showed that the Omicron VOC is more stable than the ancestral strain on surfaces (37, 38). In the present study, we prepared the inocula under identical conditions and mixed them with biologically relevant body fluids to calculate the virus decay rates over a period of time under different environmental conditions. In contrast, the inocula were prepared in virus transport medium (19, 37) or PBS following ultracentrifugation (38), and 2 or 5 µL of inoculum was placed on surfaces in previous studies. These differences in methodology and data analysis might explain why our results indicate that the ancestral lineage A strain was more stable than the VOCs used this study.

There are several limitations in the present study. First, we prepared the virus stocks from a Vero-TMPRSS2 cell line originating from a non-human primate. A previous study showed that influenza viruses sharing the same genetic background exhibited different virus decay rates in water based on the cell line in which they were cultivated, suggesting an influence of host/cell origin on virus stability (39). Second, we used early isolates of Alpha and Beta VOCs (isolated in November 2020) and the Omicron VOC (isolated in November 2021). Even though SARS-CoV-2 strains are classified into defined VOCs, some VOC isolates have accumulated additional mutations in their structural proteins over time, resulting in the emergence of novel sublineages within the same VOC. In addition, we found spontaneous amino acid substitutions that emerged after serial passages in cell culture, such as Q677H and R682W in the spike protein of the Beta VOC. Amino acid substitutions from additional or spontaneous mutations may also impact virus stability.

The main route of SARS-CoV-2 transmission occurs when an individual directly inhales respiratory droplets or aerosols from a nearby infected person. On the other hand, fomite transmission seems to play a lesser role in SARS-CoV-2 transmission. Through the evolutionary processes, SARS-CoV-2 has acquired the ability to adapt and survive in various hostile environments; it was able to evade pre-existing immunity and to evolve to more efficient transmissibility. Given this scenario, it is plausible that less stable VOCs became dominant worldwide because such variants seem to show enhanced transmissibility and increased fitness in mammals. In conclusion, our data indicate that fomite transmission of SARS-cov-2 seems less likely via liquid nasal mucus and sputum. In contrast, in dried biological fluids, SARS-CoV-2 VOCs more than the ancestral strains could still pose a significant fomite transmission risk. The present work provides novel insights into the stability of SARS-CoV-2 and its VOCs and could, therefore, contribute to the development of mitigation strategies to reduce fomite transmission.

## Materials and methods

### Cell and virus

Vero-TMPRRS2 cells were cultured in Dulbecco’s Modified Eagle Medium (DMEM; Corning, Manassas, VA, USA) supplemented with 10% fetal bovine serum (FBS; R&D systems, Flower Branch, GA, USA), 1% antibiotic-antimycotic solutions (Gibco, Grand Island, NY, USA), and Geneticin (Gibco, Grand Island, NY, USA) and maintained at 37 °C in a humidified 5% CO2 incubator. In this study, we used four different SARS-CoV-2 strains; (1) USA-WA/2020 which was isolated from the first U.S. patient in January 2020 (BEI catalog number: NR-52281; herein as WA-1), (2) hCoV-19/England/204820464/2020 which was isolated in November 2020 in United Kingdom (NR-54000; herein as Alpha VOC), (3) hCoV-19/South Africa/KRISP-K005325/2020 which was isolated in November 2020 in South Africa (NR-54009; herein as Beta VOC), and (4) hCoV-19/USA/NY-MSHSPSP-PV44476/2021 which was isolated in November 2021 in New York, USA (herein as Omicron VOC). Virus stocks were prepared in Vero-TMPRRS2 cells which were maintained in virus growth medium [DMEM supplemented with 5% FBS and 1% antibiotic-antimycotic solutions]. The titer of virus stocks was determined using end-point titration in Vero-TMPRRS2 cells.

### Next-generation sequencing

Viral RNA was extracted from virus stocks using a magnetic bead based automatic extraction system (Taco™ DNA/RNA Extraction Kit, GeneReach, Lexington, MA, USA) and sequenced by next-generation sequencing using an Illumina NextSeq (Illumina, Inc., San Diego, CA, USA) as reported previously (40-42). Briefly, nucleic acid extractions were performed by combining equal amounts of cell culture supernatants with RLT Lysis Buffer (Qiagen, Germantown, MA, USA), with 200 µL of the lysate used for magnetic bead-based extraction according to the manufacturer’s protocol. SARS-CoV-2 cDNA was then synthesized and amplified using the ARTIC-V3 RT-PCR protocol (reference: Josh Quick 2020. nCOV-2019 sequencing protocol vs (GunIt), https://gx.doi.org/10.17504/protocols.io.bdp7i5rn, followed by library preparation using a Nextera XT library prep kit (Illumina, Inc., San Diego, CA, USA) according to manufacturer’s protocols. The libraries were then sequenced with the Illumina NextSeq platform using paired-end 150 bp reads. Reads were then demultiplexed and parsed into individual sample files that were imported into CLC Genomics Workbench version 7.5 (Qiagen, Germantown, MD, USA) for analysis. Reads were trimmed to remove ambiguous nucleotides at the 5’ end and filtered to remove low quality and short reads. To determine amino acid substitutions in the Alpha and Beta VOCs, sequencing reads were mapped to the WA-1 (GISAID accession ID: EPI_ISL_404895) consensus sequence, followed by analysis with the low frequency variant detector program in CLC Genomics workbench to determine non-synonymous substitutions. The consensus sequences for each of the variant isolates were extracted from the read mappings. Consensus sequences were then aligned with published sequences from the GISAID database (GISAID accession ID: EPI_ISL_683466 for the Alpha VOC, EPI_ISL_678615 for the Beta VOC, and EPI_ISL_7908052 for Omicron VOC) to manually inspect all identified mutations. The undetermined regions in next-generation sequencing were further confirmed by Sanger sequencing using primers in the Midnight protocol (dx.doi.org/10.17504/protocols.io.bwyppfvn).

### Virus stability assay

To prevent potential cross-contamination during the work, the stability assay for each virus strain was performed at separate times, and surfaces of the biosafety cabinet and equipment were thoroughly decontaminated between work with different strains using appropriate disinfectant. In the first study, inoculums of WA-1, Alpha VOC, and Beta VOC were prepared by diluting the virus stock in virus growth medium at the concentration of 10^7^ TCID_50_/mL. A total 5 μL of the diluted virus (5 × 10^4^ TCID_50_) was directly mixed with 0.1 g to 0.2 g of human nasal mucus or sputum (Lee Biosolutions Inc., Maryland Heights, MO, USA) in a 2 mL tube or on a stainless steel surface in a 12-well plate. The same amount of the virus was mixed with 45 µL of human saliva (Lee Biosolutions) or medium and transferred into a 2 mL tube or onto stainless steel in the 12-well plate. The mixture on the steel surface was completely air-dried in a biosafety cabinet for 4 hours. The virus-spiked biologicals in the 2 mL tube and on stainless steel were then placed in a temperature- and humidity-controlled chamber (Nor-Lake Scientific, Hudson, WI, USA) under four different environmental conditions: 21 °C/60% relative humidity (RH), 25 °C/70% RH, 13 °C/66% RH, and 5 °C/75% RH. These conditions simulated the indoor, summer, spring/fall, and winter climatic conditions for the Midwestern US (15, 20). Three virus-spiked biological samples per each time point (Supplementary table 2) were subject to virus isolation in Vero-TMPRSS2 cells. Briefly, the infectious virus was recovered in 2 mL of the virus growth medium, vortexed thoroughly and filtered through a 0.45 μm syringe filter. Ten-fold serial dilutions were prepared and transferred onto Vero-TMPRSS2 cells in a 96-well plate. The presence of cytopathic effect was recorded at 4 days, and the virus titer was calculated using Reed-Muench method.

The second study was carried out to determine the stability of Omicron VOC in liquid nasal mucus and sputum. The inoculums of WA-1 and Omicron VOC were prepared in virus growth medium at the concentration of 6.2 × 10^6^ TCID_50_/mL, and 5 μL of the diluted virus (3.1 × 10^4^ TCID_50_) was directly mixed with 0.1 g to 0.2 g of human nasal mucus or sputum. The mixtures were incubated under spring/fall conditions, and samples at each time points were processed as mentioned above.

The log-transformed virus titers from the first time point (1.5 or 4 hours) to the last positive time point when at least one out of three replicates was positive were used to estimate a simple linear regression using GraphPad Prism 9 software (GraphPad, San Diego, CA, USA). The half-life value was calculated as − log_10_ (2)/slope. Statistical difference in half-life values between WA-1, Alpha VOC, and Beta VOC were evaluated by one-way analysis of variance (ANOVA), followed by multiple pairwise comparisons using Tukey’s adjustment according to the software’s instruction. In addition, statistical difference between WA-1 and Omicron VOC was tested using default analysis, which is compatible to analysis of co-variance in GraphPad Prism 9.

## Acknowledgments

We gratefully thank Velmurugan Balaraman for preparing the virus stock of WA-1 strain, Dr. Adolfo García-Sastre at Icahn School of Medicine at Mount Sinai, New York for providing two isolates, NR-54000 and NR-54009, and Dr. Michael Schotsaert at Icahn School of Medicine at Mount Sinai, New York for providing the hCoV-19/USA/NY-MSHSPSP-PV44476/2021 isolate. Funding for this study was provided through grants from the National Bio and Agro-Defense Facility (NBAF) Transition Fund from the State of Kansas, the MCB Core of the Center on Emerging and Zoonotic Infectious Diseases (CEZID) of the National Institutes of General Medical Sciences under award number P20GM130448, the NIAID Centers of Excellence for Influenza Research and Surveillance under contract number HHSN 272201400006Ct and the NIAID supported Center of Excellence for Influenza Research and Response (CEIRR) under contract number 75N93021C00016. The following reagents were deposited by the Centers for Disease Control and Prevention and obtained through BEI Resources, NIAID, NIH: SARS-Related Coronavirus 2, Isolate USA-WA1/2020, NR-52281, isolate hCoV-19/England/204820464/2020, NR-54000, contributed by Bassam Hallis, and isolate hCoV-19/South Africa/KRISP-K005325/2020, NR-54009, contributed by Alex Sigal and Tulio de Oliveira. The Omicron variant, hCoV-19/USA/NY-MSHSPSP-PV44476/2021, was originally obtained through Dr. Viviana Simon in Mount Sinai Pathogen Surveillance program.

